# Physical structure and interstitial flows govern microbial life in microenvironments

**DOI:** 10.1101/2023.09.19.558408

**Authors:** Rachel Shen, Benedict Borer, Davide Ciccarese, M. Mehdi Salek, Andrew R. Babbin

## Abstract

Most microbial life on Earth is found in localized microenvironments that collectively exert a crucial role in maintaining ecosystem health and influencing global biogeochemical cycles. In many habitats such as biofilms in aquatic systems, bacterial flocs in activated sludge, periphyton mats, or particles sinking in the ocean, these microenvironments experience sporadic or continuous flow. Depending on their microscale structure, pores and channels through the microenvironments permit localized flow that shifts the relative importance of diffusive and advective mass transport. How this flow alters nutrient supply, facilitates waste removal, drives the emergence of different microbial niches, and impacts the overall function of the microenvironments remains unclear. Here, we quantify how pores through microenvironments that permit flow can elevate nutrient supply to the resident bacterial community using a microfluidic experimental system and gain further insights from coupled population-based and computational fluid dynamics simulations. We find that the microscale structure determines the relative contribution of advection versus diffusion, and even a modest flow through a pore in the range of 10 µm s^−1^ can increase the carrying capacity of a microenvironment by 10%. Recognizing the fundamental role that microbial hotspots play in the Earth system, developing frameworks that predict how their heterogeneous morphology and potential interstitial flows change microbial function and collectively alter global scale fluxes is critical.

## Introduction

Microbial life in natural and engineered environments most often exists in discretely localized microenvironments or hotspots such as biofilms, aggregates, flocs, particles, and granules, to name a few^1^. Unifying these environments is the principle of living in confinement, whereby the collective activity of the community creates steep nutrient and oxygen gradients within and around these environments. In turn, the resulting diffusive mass transport dictates nutrient acquisition of individual cells and gives rise to a landscape of different metabolisms, all across a spatial scale of a few micrometers. These self-engineered microenvironments are not isolated occurrences in specific habitats but are abundant, ubiquitous, and collectively play a crucial role in driving global biogeochemistry^2^. For instance, soil bacterial hotspots such as aggregates and the rhizosphere^3^ are important sources of greenhouse gases^4^ due to the localized development of anoxic niches^5^. In the oceans, similar anoxic microenvironments are observed in settling marine snow^6,7^ that support anaerobic metabolisms^8^ and can nearly double the niche size for marine denitrification on a global scale^9^. The occurrence of anoxic microenvironments due to diffusive limitations is not exclusive to natural environments. They also emerge in systems that are optimized for oxygen exchange such as flocs in activated sludge^10^ or within pulmonary biofilms of cystic fibrosis patients^11^. At finer scales within a microenvironment, the steep nutrient gradients govern the spatial organization of the bacterial community^12,13^ where even a single species shows phenotypic heterogeneity depending on the precise location of the cells inside the hotspot^13–15^. This diffusive limitation not only generates anoxic conditions but also affects all resources when localized consumption exceeds supply.

Yet, typically overlooked is the role of advection in addition to diffusion in setting the chemical landscape where flow around microbial assemblages can impact the way organisms perceive nutrient gradients and alter nutrient fluxes^16^. For instance, the degradation rate of alginate particles increases due to enhanced mass fluxes across the particle as a function of flow velocity^17^. These relationships between the localized flow and the increase in mass transport due to bulk advection are primarily studied for simplified geometries such as solid spheres or cylinders. Natural microbial hotspots that encounter flow such as marine particles^18^, biofilms in streams and intertidal zones^19–21^ or flocs in activated sludge^22^ typically manifest irregular shapes and structures that may support interstitial flow through pores or gaps between cell clusters. The relative importance of advective versus diffusive nutrient transport in these systems is then a function of the diffusivity of the nutrient, the localized flow velocity, and the spatial scale^16^. Mathematically, this is captured by the dimensionless Péclet number (Pe), defined as the ratio between the rate of advective and diffusive mass transport (Eqn. 1) where L is the spatial scale (e.g., the diameter of the cylinder or sphere), v the flow velocity and D the diffusivity of the considered nutrient.

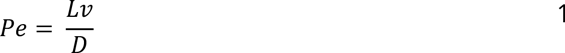

At Pe smaller than unity, nutrients are primarily transported by molecular diffusion whereas advective transport dominates at high Pe. Since diffusive fluxes are restricted to short distances, a landscape of Pe emerges at the microscale^23^ with regions of high advective mass transport due to localized fluid flow and regions where mass transport is restricted to diffusive fluxes. For small molecules (diffusivities on the order of 10^−9^ m^2^ s^−1^), a pore of radius 50 µm and flow velocity of 10 µm s^−1^ results in a Pe of 1, dimensions which are relevant to many microbial hotspots. This suggests that hotspots may often experience tipping points between diffusive and advective nutrient supply. Recently, the impact of the microscale structure on the localized flow around marine particles has been investigated^24^, but the overall effect of the microenvironment morphology on nutrient supply, microbial growth conditions within the porous structure, and the carrying capacity of individual microenvironments remain unclear.

In the present study, we aim to understand how the interstitial flow through a microbial microenvironment alters the spatial growth conditions, and how these alterations affect the overall carrying capacity of the microbial hotspots. We hypothesize that large pores that support interstitial flow can critically elevate nutrient supply to the microenvironment with significant consequences for microbial growth and community function. The objectives of this study are (i) to quantify the influence of advective nutrient supply at the microscale on the growth of bacterial populations and (ii) to generalize these insights using simple engineering and fluid mechanics principles that transcend phenomenological-specific examples. Understanding the bidirectional interaction between bacterial populations and their heterogeneous microenvironment is fundamental to quantifying the influence of such microenvironments on global-scale biogeochemical fluxes and developing predictive capabilities for engineering applications such as in wastewater treatment plants or bioremediation.

## Results

### Bacterial carrying capacity as a function of interstitial flow

We quantify the relationship between flow through microenvironments, the elevated nutrient supply, and the proliferation of the resident bacterial population by observing the growth of bacterial colonies in hydrogel particles subjected to flow in millifluidic devices (Fig. 1a). We control the bulk flow velocity (115 µm s^−1^) around 3 mm diameter agarose hydrogel particles and create a channel through the particles of approximately 400 µm width to imitate a large pore (Fig. 1b). The contribution of advective fluxes to the total nutrient supply is controlled by changing the angle of this channel (0°, 30°, 60°, 90°) relative to the flow direction (Fig. 1c and 1d, see Methods section for details on the device and particle fabrication). We compare these particles to a solid control particle (i.e., without a channel) that does not permit any interstitial flows. We determine the influence of interstitial flow on the particle carrying capacity by examining the pattern of colony size as a function of the distance from the nutrient-rich particle periphery in solid particles (that are restricted to diffusive fluxes) and ascribe large colonies that diverge from this pattern in particles containing a channel to advective fluxes. We quantify the increase in biomass due to advection as the ratio of the total observed biomass within those particles to a hypothetical scenario where we omit the anomalous colonies (for more details, see Supplementary Fig. 1 and 2). The biomass fold-increases calculated here and reported throughout the rest of the manuscript are thus an increase in biomass within the particles due to advective nutrient contributions relative to a solid particle. This approach effectively normalizes the biological variance between replicates and isolates the influence of elevated nutrient supply due to advective fluxes (Fig. 2a). There is a significant influence of the channel angle (as a proxy for increased nutrient flux) on the relative biomass increase (Welch ANOVA, p-value = 0.0012) with the highest relative increase for the in-line channel (0° angle, mean ± std relative increase of 7.52% ± 4.08%). This influence decreases with increasing channel angle where a perpendicular channel to the flow (90° angle, mean ± std relative increase of 1.18% ± 1.33%) is indistinguishable from a solid particle (mean ± std relative increase of 1.38% ± 1.45%). Solid particles show a slight relative biomass increase due to outliers (i.e., individual large colonies deep inside of the particles that are outliers when compared to the general trend, Supplementary Fig. 1 and 2, Supplementary Table 1).

**Figure 1:**
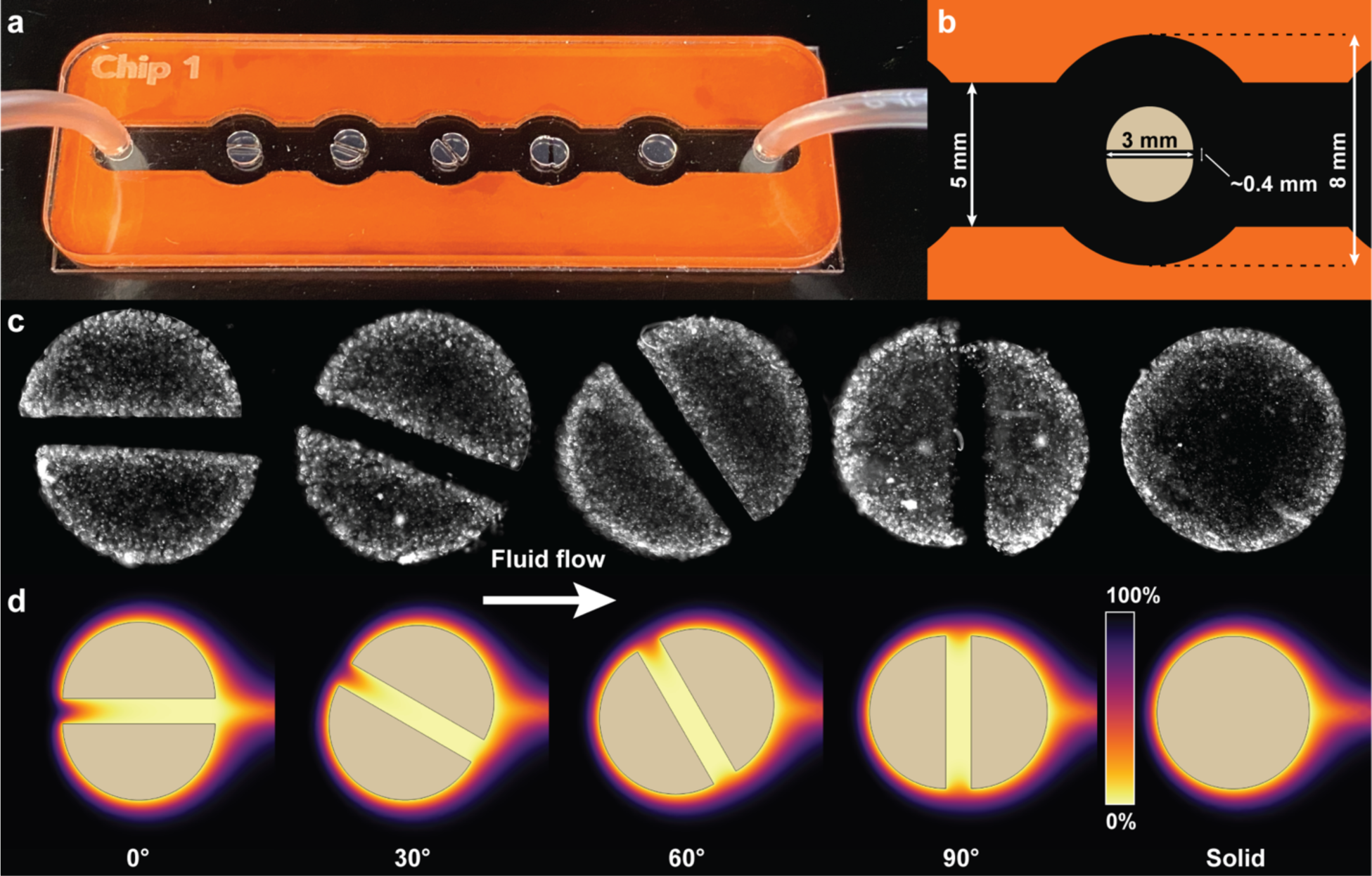
Visualization of the experimental system and predicted oxygen concentrations. **a)** Photograph of the experimental system highlighting the flow chamber and hydrogel particles with different angular orientations of the channels. **b)** Dimensions of the flow chamber and particles. **c)** Micrograph of example particles showing bacterial colonies throughout the hydrogel particles after 24h incubation. Brighter areas correspond to higher bacterial density. The contrast in these images has been increased evenly to highlight bacterial colonies. **d)** Spatial oxygen saturation around the hydrogel particles after 24h predicted with COMSOL.

**Figure 2:**
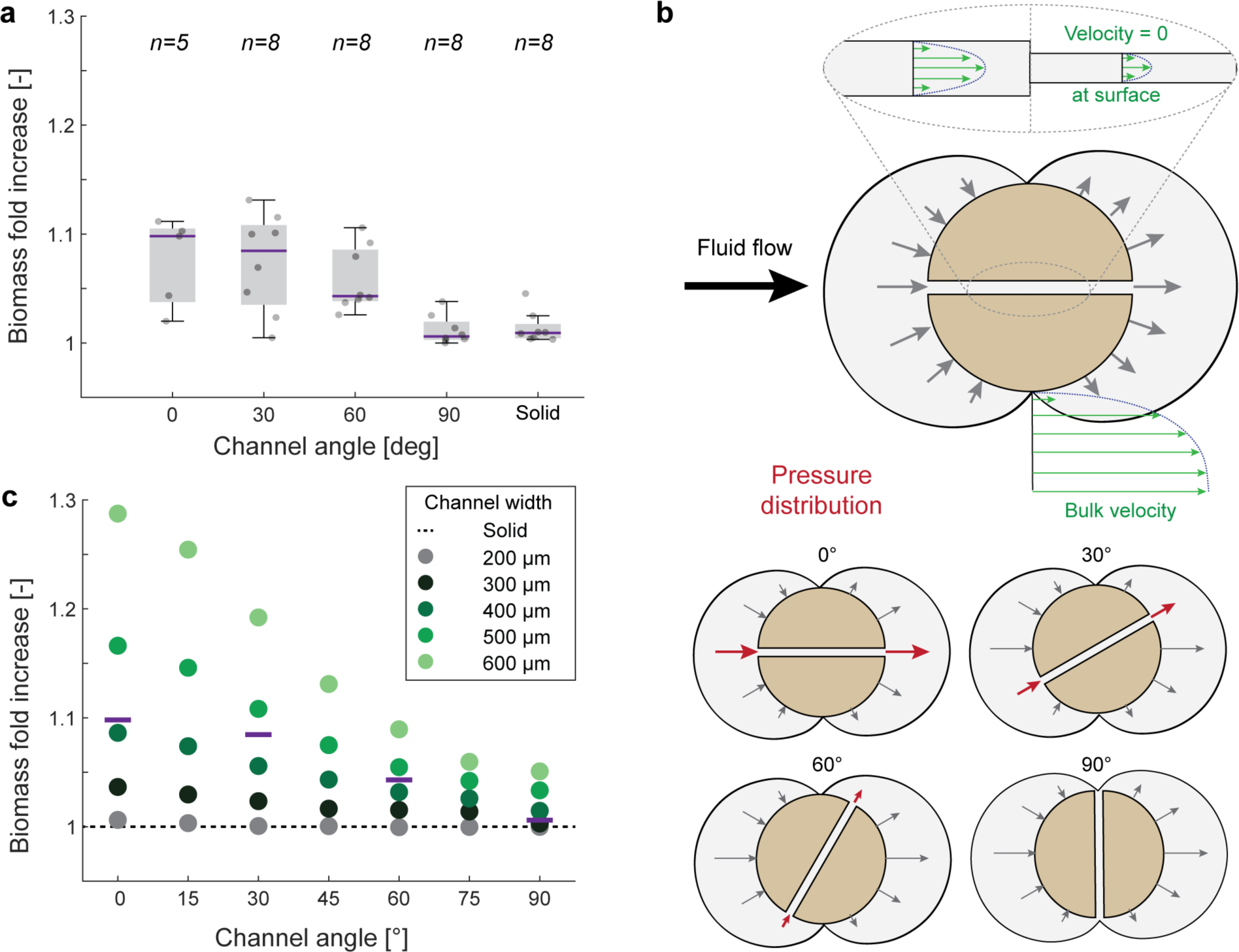
Relative biomass increase as a function of channel angle. **a)** Relative biomass increase as a function of the channel angle where n is the number of experimental replicates. The relative increase is calculated as the increase in biomass when compared to a solid particle as described in the methods section. **b)** Distribution of pressure and flow velocity around and within a cylinder containing a channel. The pressure envelope (shown in gray color) conceptually shows the distribution of the pressure magnitude around the cylinder whereas the gray arrows indicate the directionality and approximate magnitude of the pressure. The orientation of the channel with respect to the bulk flow direction further determines the pressure gradient along the channel axis. Green arrows indicate the flow velocity around the particle and within the channel. Narrow channels result in more frictional resistance and thus a lower flow velocity when compared to wider channels at the same pressure gradient along the channel axis. **c)** Relative biomasses increase as a function of the channel angle and channel widths when simulated using COMSOL. Purple lines are the mean relative biomass increase from the experimental data with an approximate channel width of 400 µm.

We hypothesize that the increase in growth observed in the presence of channels is due to elevated nutrient supply from advective fluxes. Laminar flow, a flow regime that is characterized by discrete water parcels following smooth paths and the absence of turbulent mixing, around a cylinder or sphere results in a positive relative pressure (relative to the bulk fluid pressure) at the surface facing the flow and negative relative pressure on the surface leeward of the flow (Fig. 2b). The resulting pressure gradient along the axis of the channel accelerates the fluid. Within the channel, fluid that interfaces with the channel walls has a velocity of 0 (no-slip assumption). Attraction between individual water molecules creates a drag force between adjacent fluid layers which results in a frictional resistance that decelerates the fluid. This frictional resistance is proportional to the flow velocity in the channel and inversely proportional to the channel size (i.e., narrower channels result in higher frictional forces and lower flow velocity compared to wider channels exposed to the same pressure gradient along the channel axis). The steady-state flow velocity inside the channel is thus determined when acceleration due to the pressure gradient force equals deceleration due to frictional resistance. The orientation of the channel inside the cylinder governs the pressure gradient across the channel due to the difference in pressure around the particle. A channel that is in line with the flow results in the largest pressure gradient (and thus the highest flow through the channel) whereas a channel perpendicular to the flow results in a negligible pressure gradient (and thus no corresponding flow). A spatial visualization of the flow magnitude and direction is shown in Fig. 3 for a bulk flow velocity of 115.5 µm s^−1^ and channel angles of 0°, 45°, and 90°. To summarize, the flow through a channel is determined by the bulk fluid velocity, channel width, and orientation with respect to the bulk flow direction.

**Figure 3:**
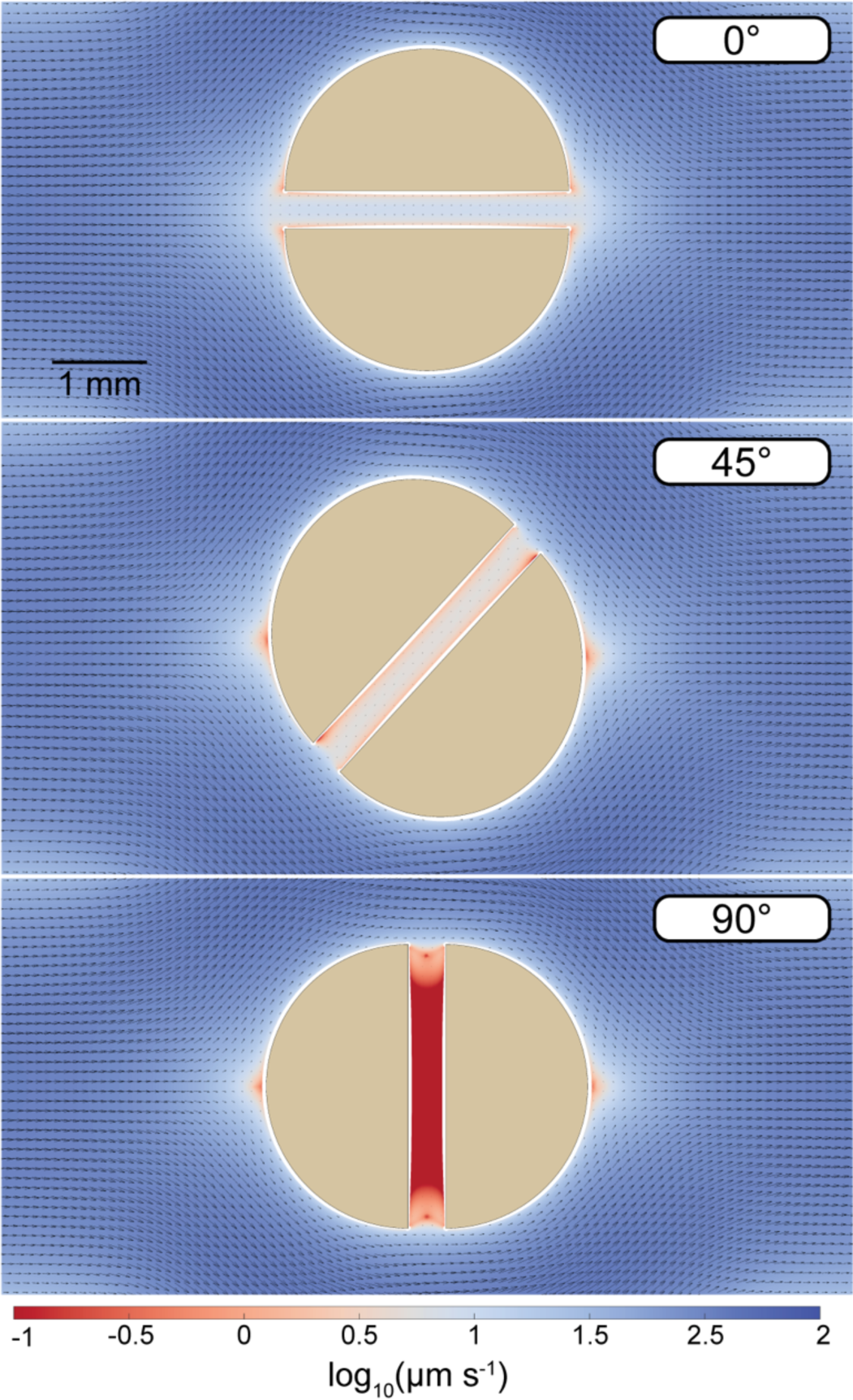
Resulting velocity field around and through the particles as a function of the channel angle. For a channel in line with the flow direction (0° angle), the stagnation point upstream of the particle (the location where the streamline directly intercepts the particle) disappears since flow through the approaching flow passes through the channel. A channel at an angle of 45° to the approaching flow results in flow through the channel albeit at a lower magnitude compared to the in-line channel. If the channel is perpendicular to the flow (90° angle), there is no pressure difference across the channel and the resulting net flow through the channel is thus 0.

To gain insights into how the microscale geometry and associated advective fluxes drive bacterial growth, we adopted a digital twin model created in the Multiphysics modeling software COMSOL (details of the model can be found in the Methods section). The model predicts a comparable relative biomass increase for particles with a channel width of 400 µm (congruent to the experiments, Fig. 2c). We further used the model to explore the influence of different channel widths (200 µm to 600 µm) and flow velocities (ranging from 11.5 µm s^−1^ to 1150 µm s^−1^, Supplementary Fig. 3). For all flow velocities and channel widths, we additionally explored intermediate angles compared to the physical experiments (15°, 45°, 75°). However, we focus on the same angles (0°, 30°, 60°, 90°) as in the experiments for visualization due to the predictable behavior of the intermediate angles (Fig. 2c). At higher bulk flow velocities (simulated as 1150 µm s^−1^, or approximately 100 m d^−1^) even a small channel or pore (300 µm) with a pore flow velocity of approximately 50 µm s^−1^ can increase the total population growth by more than 12% (Supplementary Table 2).

We further explore the influence of different flow velocities and pore characteristics (channel angle and width) on the relative biomass increase by quantifying the total flux of the limiting substrate (in our case oxygen) to the particles in the simulations (Fig. 4). Oxygen flux via the channel is the main driver of the relative biomass increase (Fig. 4a). The relative channel flux contribution is the ratio of the oxygen flux through the channel walls to the total oxygen flux (i.e., the sum of flux from the channel and particle surface). For wide channels at high flow velocities, oxygen supply through the channel can contribute up to 35% of the total supply to the particle. Interestingly, this is below the relative contribution of the channel surface to the total surface (39% calculated based on geometry), suggesting that even at 600 µm channel width, the two halves do not yet behave like distinct particles from a nutrient supply perspective. Contrary to our initial expectation, the fluid collection efficiency (defined as the fraction of liquid passing through the particle from the total approaching liquid) is not a good predictor of the relative biomass increase (Fig. 4b). This is because the fraction of fluid passing through the channel from the total fluid approaching the particle does not change with bulk flow velocity and therefore does not reflect the elevated nutrient supply and relative biomass increase through the channel (Supplementary Fig. 4). As shown in Fig. 4b, even though particles show very similar fluid collection efficiencies for larger channels at different bulk flow velocities (∼2%), the relative biomass increase is vastly different, ranging from ∼10% for fluid velocities of 11.5 µm s^−1^ to ∼50% for fluid velocities of 1150 µm s^−1^.

**Figure 4:**
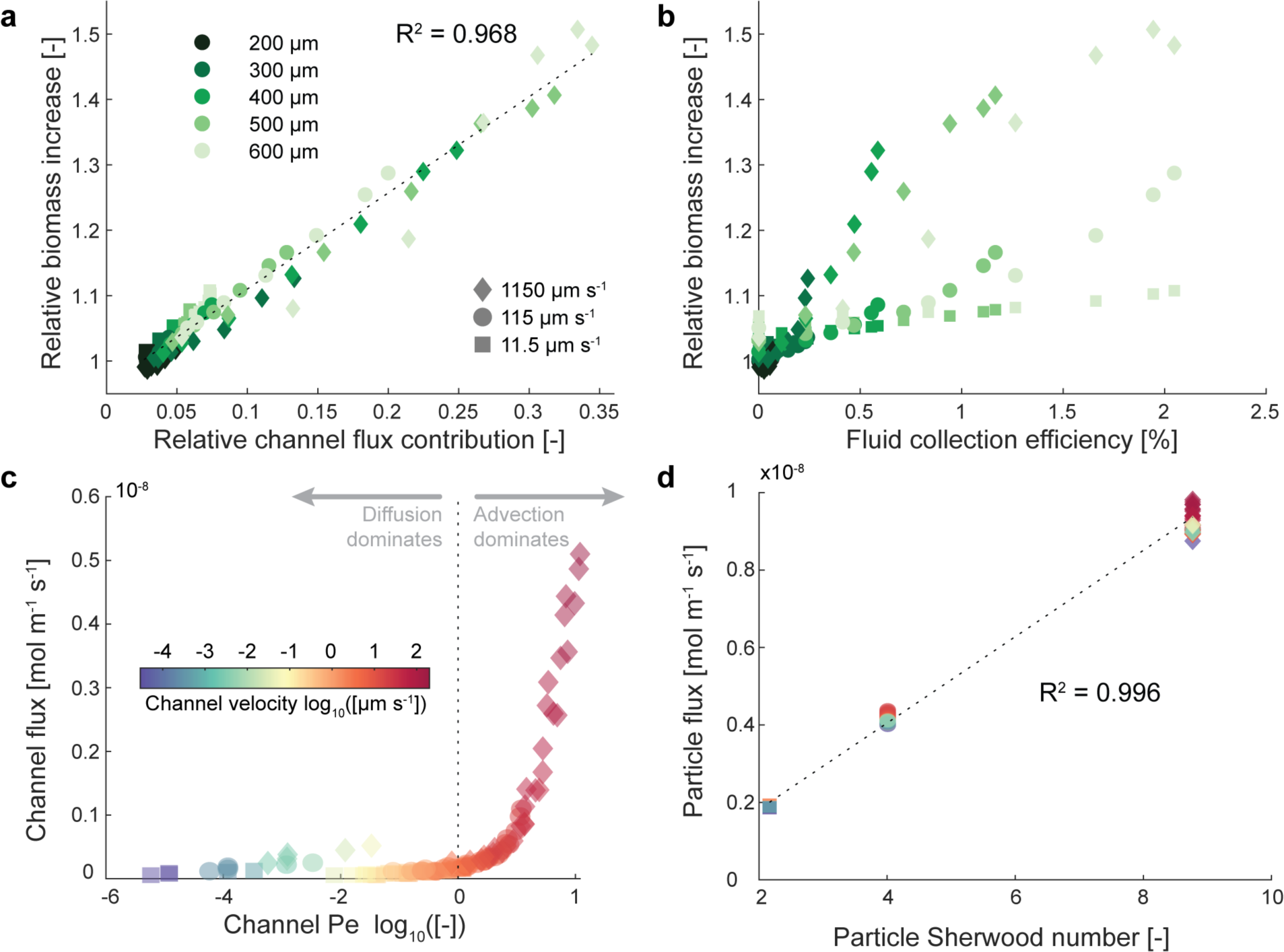
Contribution of channel and particle flux to the relative biomass increase. **a)** Relation between the relative channel flux contribution (percentage of the total flux through the channel wall) to the relative biomass increase. **b)** Relation between the fluid collection efficiency (the fraction of the flow passing through the particle from the flow approaching the particle) to the relative biomass increase. **c)** Relation between the channel Pe and the total flux through the channel. **d)** Relation between the particle Sherwood number and the total flux across the particle surface (neglecting flux through the channel).

To disentangle the relative contribution by advective flows through the channels and increase in nutrient flux due to flow velocity, we calculate the Péclet number (Pe) associated with the channel. As shown in Fig. 4c, Pe is a good predictor of how advective fluxes enhance the total nutrient flux through the channel. Overall, nutrient flux through the channel is only significant at Pe > 1 where advective mass transport dominates (Fig. 4c). However, since Pe is a function of the fluid velocity inside the channel (in our case driven by the bulk fluid velocity, angle of the channel with respect to the bulk fluid flow, and width of the channel) and channel width, different combinations of these parameters give rise to a similar regime when it comes to the relative importance of diffusive versus advective fluxes. For instance, channel widths of 200 µm, 400 µm, and 600 µm at angles of 75°, 75°, and 45° and bulk flow velocities of 1150 µm s^−1^, 115 µm s^−1^, and 11.5 µm s^−1^, result in a very similar Pe of 0.8, 0.7, and 0.7, respectively. This illustrates how vastly different bulk flow regimes and microscale geometries give rise to comparable nutrient supply and growth conditions from a microbial perspective. In addition to the elevated nutrient supply through the channel, a higher bulk flow velocity also increases the nutrient flux across the particle’s hemispherical surface (Fig. 4d). This is due to a thinning in the diffusive boundary layer which is captured by the dimensionless Sherwood number (defined as the ratio of the total combined mass transport from diffusive and advective fluxes to diffusive mass transport alone^25^). For our system, we can calculate the Sherwood number from the cylinder diameter, bulk flow velocity, and nutrient diffusivity^25^ which results in Sherwood numbers of 8.7, 4 and 2.2 for bulk flow velocities of 1150 µm s^−1^, 115 µm s^−1^, and 11.5 µm s^−1^, respectively. These correlate well with the overall increase in nutrient flux through the hemispherical surface as shown in Fig. 4d. Importantly, a Sherwood number of 2.2 suggests that despite an overall low bulk flow velocity of 11.5 µm s^−1^, the additional advective flux due to flow around the cylinder more than doubles the overall nutrient supply to the resident microbial community even in the absence of any channel or pores through the cylinder.

### Microscale spatial patterns emerge from localized nutrient availability

In addition to affecting the overall microbial community in the particles, emerging nutrient gradients change the localized growth conditions at finer scales within the particle (Fig. 5). In the experiments, we see a change in the observed colony size as a function of the distance of the channel entrance. Here and throughout the manuscript, the distance from the channel entrance is the physical distance from the channel entrance that is exposed to the oncoming flow (“windward”). Bacterial colonies show a symmetrical pattern for the 90° angled channel with larger colonies at the channel periphery (0 mm and 3 mm) and smaller colonies at the center of the particle (distance from the channel entrance of ∼1.5 mm, Fig. 5a). In this case, there is no flow through the channel and the increase in colony size at the periphery is due to an increase in diffusive fluxes from the surrounding flow. With increasing flow through the channel from 60° to 0° (in line with the flow), this pattern becomes more skewed toward larger colonies directly at the channel entrance. This pattern is congruent to the increased oxygen availability supported by advective fluxes into the particle (Fig. 5b). Channels perpendicular to the flow show a very rapid decline in oxygen concentration along the channel axis whereas oxygen may penetrate deeply into the particle for channels in-line with the flow. The oxygen penetration depth (defined as the distance from the channel entrance when the oxygen concentration falls below 1 µmol L^−1^) is critically dependent on the flow within the channel (driven by the channel orientation and width) and the bulk flow velocity (see Supplementary Table 3). 1 µmol L^−1^ oxygen was taken as a semi-arbitrary threshold where aerobic respiration becomes negligible^26^. For the case of a channel perpendicular to flow, oxygen penetration is independent of the bulk fluid velocity (e.g., 450 µm into a 400 µm wide channel at all velocities), suggesting that oxygen penetration is solely from diffusion. In contrast, the penetration depth for a 400 µm channel at a 30° angle relative to the bulk flow direction increases from 450 µm to 750 µm to complete penetration (>3mm) for flow velocities of 11.5 µm s^−1^, 115 µm s^−1^, and 1150 µm s^−1^, respectively. For a high flow velocity of 1150 µm s^−1^, even a narrower channel of 300 µm increases the oxygen penetration depth nearly eightfold from 340 µm (when the channel is perpendicular to the flow direction and restricted to diffusive penetration) to 2.5 mm when the channel is in line with the flow.

**Figure 5:**
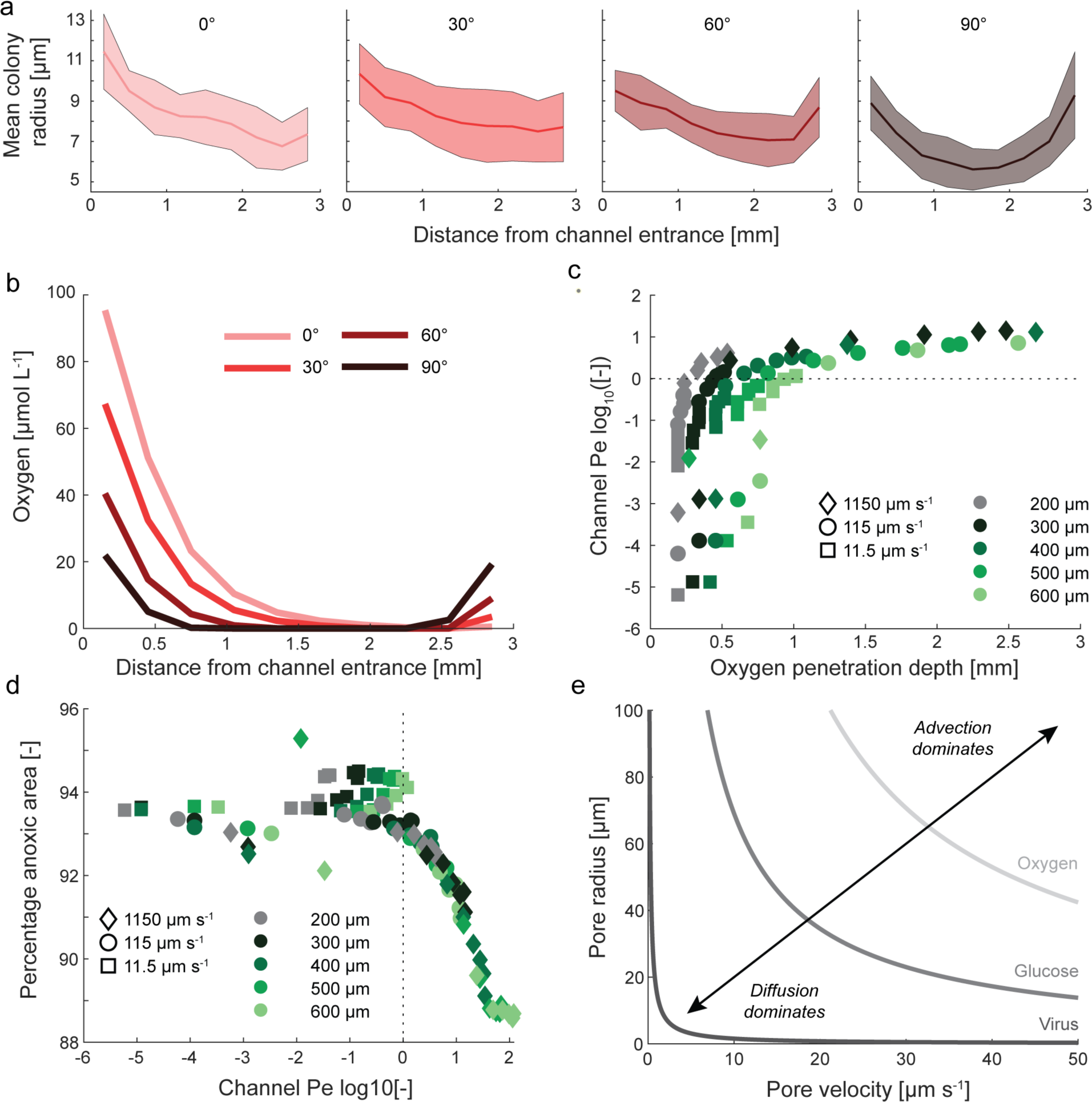
Microscale spatial patterns of colony size and oxygen distribution within the particles. **a)** Colony size as a function of the distance to the channel entrance (channel entrance exposed to oncoming flow) for all channel angles in the experiments. Solid lines show the mean with the shaded area ± one SD around the mean. **b)** Oxygen concentration at the channel center as a function of the distance to the channel entrance for a flow velocity of 115 µm s^−1^ and 400 µm channel width. **c)** Oxygen penetration depth (defined as the distance from the channel entrance when oxygen at the channel center drops below 1 µmol L^−1^) in relation to the channel Pe for all flow velocities, channel widths, and angles. Penetration depths that exceed 3 mm are omitted. **d)** Percentage of the area that is anoxic (defined as the percentage of the area where oxygen concentrations are below 1 µmol L^−1^) in relation to the channel Pe for all flow velocities, channel widths, and angles. **e)** Delineation between advective and diffusive dominated transport (curves indicate the threshold with Pe = 1) considering nutrients and particles with varying diffusivities.

The rapid and non-linear increase of the penetration depth with flow velocity can be explained by the channel Pe number (Fig. 5c). At lower Pe (values below the dashed line in Fig. 5c), the channel width is an important factor driving the penetration depth since the total diffusive oxygen flux into a wide channel is greater when compared to a thin channel (proportional to the channel cross-sectional area). Advective fluxes on the other hand dominate at higher Pe where oxygen can penetrate the whole particle, especially at higher flow velocities and wide channels. Channels, where oxygen does not fall below 1 µmol L^−1^ (i.e., oxygen fully penetrates the channel), have a mean Pe of 50. Advective fluxes not only determine the increase in growth but also affect the potential metabolisms occurring within the microbial hotspots. Although most of the particles remain anoxic in all conditions, an increase in oxygen supply via advective fluxes reduces the percentage of the area within the particle that is anoxic (Fig. 5d). When diffusion dominates (Pe < 1, to the left of the dotted line in Fig. 5d), the percentage of the particle that experiences oxygen levels above 1 µmol L^−1^ is approximately 6%. With the onset of significant advective fluxes (Pe > 1, to the right of the dotted line in Fig. 5d), the oxic area rapidly increases to approximately 12%. Although this may represent a minor increase relative to the total particle area, it doubles the niche where microbial cells can maintain aerobic respiration.

Finally, advective fluxes not only enhance oxygen supply but influence all mass transport to and from the particle. Using the Péclet number, we can compute the pore characteristics in which advective fluxes become significant for different entities (Fig. 5e). Here, the shape of the curves (which indicate Pe = 1) is purely a function of the entity’s diffusivity and separates pore characteristics where diffusion dominates (combinations of flow velocity and pore size below the curve) from those where advective fluxes dominate mass transport. For rapidly diffusing entities such as oxygen, diffusive fluxes dominate except for large pores with high flow velocities. For slower diffusing entities such as glucose or especially small particles such as viruses, small interstitial flow rates in narrow pores already contribute significantly to the overall transport of the entity through advective fluxes. As such, limitation by larger substrates can be greatly reduced, and viral infection rates greatly enhanced by the intra-particle advective flow.

## Discussion

Our results highlight the importance of interstitial advection for nutrient supply and the proliferation of bacteria in microenvironments that are exposed to bulk flow such as biofilms on riverbeds and intertidal zones, the periphyton, marine snow particles, or flocs in activated sludge. We controlled the pore velocity (and associated advective nutrient supply) by changing the angle relative to the flow to achieve a wide range of conditions without changing the particle’s geometry. By doing so, we strategically disentangled the impact of elevated nutrient supply from interstitial advective fluxes through the channels (Fig. 4c) versus the particle surface related to the bulk flow velocity (Fig. 4d). In our system, a modest pore velocity of 10 µm s^−1^ supported an increase in carrying capacity of approximately 10%, where faster flow velocities inside channels support an increase in carrying capacity of up to 50% (Supplementary Tables 2 and 4). Considering that we find these results for a rapidly diffusing substrate such as oxygen, advective fluxes may contribute a substantial fraction of the total nutrient supply to microenvironments for substrates with lower diffusivities such as larger organic molecules or even viruses (Fig. 5e).

The question remains if pores in microenvironments that support advective fluxes are widespread or if exopolymers produced by the resident microbial community will rapidly fill any voids within the microenvironments^27^. Early studies on flow through biofilms^28^ revealed a network of pores in between bacterial clusters of up to 50 µm radius that support high interstitial flows between 2000 and 15000 µm s^−1^. Similarly, visualizing the flow field around and through bacterial flocs^29^ and marine aggregates^30^ using particle image velocimetry have demonstrated the existence of flow penetrating through the particles, although similar experiments investigating the micro-hydrodynamics around marine snow particles did not show conclusive evidence supporting interstitial flow^24^. However, in these latter experiments, the marine particles were impaled on a needle to visualize the flow which selected for more robust and potentially less porous particles. Natural marine particles also demonstrate high fluid collection efficiencies (i.e., the percentage of the approaching flow that passes through the particles) ranging from 12% to 40% for highly porous particles^30^, which is an order of magnitude higher when compared to the collection efficiency calculated in our experiments (Fig. 4b) and suggests that the carrying capacity in the former may be elevated beyond our observations. Finally, it has been shown experimentally by comparing measured oxygen consumption rates and calculated oxygen supply from oxygen gradients that diffusion alone cannot supply sufficient oxygen to a marine particle to explain its high consumption^31^. Within those experiments, additional interstitial advective fluxes between 5–40 µm s^−1^ were required to account for the discrepancy between observed oxygen consumption and that predicted solely by diffusive theory^31^. Together, these experimental results suggest that large pores and pore networks through particles and within biofilms that permit advective fluxes are omnipresent features that greatly alter the localized growth conditions of microbial cells within the microenvironment.

Within an individual microenvironment, multiple gradients emerge rapidly as a function of bacterial nutrient consumption or intermediate metabolite production. In diffusion-dominated microenvironments, localized anoxia develops^32^ that permits anoxic metabolisms such as denitrification^9,13,33^ or sulfate reduction^10^. Enhanced oxygen supply due to advective fluxes can change the landscape of metabolisms occurring within the microenvironment and restrict anoxic metabolisms to a fraction of the microenvironment when compared to a diffusion-dominated scenario. An oxygenated periphery can significantly reduce the anoxic niche in the presence of a channel that elevates oxygen supply when compared to a solid particle (Fig. 5d), especially when considering that nutrients supplied from the periphery are now intercepted in the oxygenated layer. Fundamentally, this is only relevant for large particles with high interstitial flows that support deep oxygen penetration into the channel via advective fluxes (Fig. 5c). In contrast, we observe a shallower oxygen penetration depth into the channel for most conditions. This presents an interesting scenario where advective fluxes do occur, but microbes keep pace (they grow larger) and oxygen is rapidly consumed such that the bulk of the particle remains anoxic. These advective fluxes may thus increase the supply of anaerobically respired nutrients such as nitrate or sulfate without disrupting the anoxic niche. By doing so, the permeation of the particles not only increases the total carrying capacity but additionally elevates anaerobic metabolism occurring within the microenvironment. This is not only the case for electron acceptors such as described above but for any nutrient supplied from the bulk liquid (Fig. 5e). Advective fluxes may shift the limiting nutrient within a microbial hotspot, especially if there is a large discrepancy in diffusivity such as between oxygen and larger organic molecules. In a purely diffusion-dominated system, substrates with higher diffusivity will result in a higher overall substrate flux for the same cross-sectional area when compared to a substrate with lower diffusivity (assuming equal concentration and driving gradient). This generates a scenario where one substrate is more available simply due to the diffusive characteristics. In contrast, if advective fluxes dominate, they are supplied at their relative concentration in the bulk fluid, and the diffusive gradient is disrupted, which can shift the system to a different limiting nutrient depending on the uptake stoichiometry between the substrates. For example, advective fluxes in a carbon-limited system may elevate the carbon supply such that the system becomes oxygen limited since their diffusivities often differ significantly. Oxygen^34^ has a diffusion coefficient of 18×10^−10^ m^2^ s^−1^ whereas L-valine^35^, citrate^36^, and sucrose^37^ have diffusion coefficients of 6.6, 5.9, and 4.8×10^−10^ m^2^ s^−1^, respectively (or 2.7, 3 and 3.7 times lower than oxygen). Considering that the stoichiometry for aerobic respiration between oxygen and a six-carbon sugar such as citrate is 6:1, a small increase in carbon flux can rapidly result in a rapid depletion of oxygen if oxygen fluxes do not increase correspondingly. However, this process is not generalizable as it is a function of the substrate concentrations, their uptake stoichiometry, and diffusivity, amongst others.

Finally, advective fluxes may not only elevate nutrient supply, but they may also flush away intermediate metabolites from the microenvironments. For example, the release of nitrous oxide, both an intermediate metabolite in denitrification and a critical greenhouse gas, may result in a local loss of energy-yielding substrates but also modulate the interaction between these microenvironments and biogeochemical cycles. On the other hand, the removal of intermediate metabolites may also benefit the resident microbial communities in the case where these inhibit growth. This has been demonstrated for solid spheres where flow assisted in removing intermediate degradation products which amplified the overall degradation rate^17^ – a process that would be further enhanced by permeation of the particle. Localized flow and the relative change in diffusive versus advective transport also modify the way organisms such as non-motile cells or viruses interact with microenvironments. Due to their large size and associated low diffusivities, interstitial fluxes on the order of a few microns per second already elevate the potential colonization and infection rate by non-motile cells and viruses, respectively (Fig. 5e). For example, porous marine snow that permits interstitial advective flows is colonized more efficiently by both motile and non-motile cells when compared to a rough or smooth surfaced particle^37^. However, while motile cells showed an average 10-fold increase in colonization for porous particles, this varied considerably for non-motile cells ranging from a negligible increase to an over 1000-fold increase. This is because non-motile cells (and by analogy viruses) rely on a direct, stochastic encounter with the particle (where the microstructure mediates the flow in and around the particles) whereas motile cells simply benefit from a reduction in flow velocity and ability to colonize through random or chemotactically guided motion^37^. Finally, the permeation of microenvironments also changes the way these interact with each other in terms of aggregation. Activated sludge in wastewater treatment plants and marine particles develop through direct collisions of smaller colloids supplemented by exopolymeric substances produced by microbes^38,39^. Similarly, biofilms in streams or intertidal zones can import colloids as evidenced by the rapid incorporation of metal nanoparticles^40^. The presence of porosity that promotes interstitial flows can shift the dynamics of these processes. Large particles that are typically more porous scavenge smaller particles at rates that rapidly exceed theoretical collision efficiencies in the absence of particle permeation^41^.

Our results demonstrate the importance of microscale variations in flow for the development and functioning of microbial communities within microenvironments due to a change in nutrient supply. Indeed, the overall supply to microbial hotspots is a delicate balance between advective and diffusive nutrient fluxes where their relative importance is captured by the dimensionless Péclet (Pe) number. Elevated nutrient fluxes due to supply through pore spaces directly impact the carrying capacity of the microbial hotspots (Fig. 4a) and are tightly linked to the Pe (Fig. 4c). The increase in nutrient supply due to dominant advective fluxes defined as Pe ≥ 1 also delineates the threshold where oxygen penetration into the channel or pore becomes significant (Fig. 5c) and alters the fraction of the hotspot where aerobic growth may occur (Fig. 5d). Importantly, the Pe can be calculated directly from the geometry of the pore spaces, localized fluid velocity, and diffusivity of the nutrient under consideration. From this perspective, characterizing the localized flow regime and microscale geometry of microbial hotspots can explain potential shifts in microbial metabolism within microbial hotspots, but also enable the informed design for engineering applications.

In summary, the elevated mass transport from advection on top of diffusion not only impacts the carrying capacity of microbial hotspots but also shapes the emerging niches within the microenvironments with consequences for the spatial occurrence of diverse metabolisms, the proliferation of specialized species, and the overall diversity of the resident community. These changes in microbial communities greatly impact metabolisms associated with global biogeochemical cycles and overall ecosystem function. Furthermore, advective fluxes change the colonization or detachment of non-motile bacteria from these microenvironments or critically facilitate the contact between microbial hotspots. However, how the permeation of microenvironments and associated changes to the local biogeochemistry and ecology of microbial hotspots collectively impact the functioning of meso- and global-scale processes in the Earth system remains unknown.

## Methods

### Bacterial strains and culture conditions

We used a fluorescently tagged *Pseudomonas aeruginosa* PA14 that constitutively expresses a yellow fluorescent protein (eyfp) for all experiments^37^. We used Lennox lysogeny broth (5 g L^−1^ NaCl, BD Life Sciences, further abbreviated LB) media for overnight cultures and in the experiment and supplemented the overnight cultures with 300 µg/ml Trimethoprim (Trp) to avoid contamination.

### Fabrication of millifluidic devices

We adapted a previously described experimental system that mimics a settling marine particle amenable to microscopy^13^. We fabricated the millifluidic devices using two 1mm thick microscope slides (VWR, Vistavision Microscope Slide, 75 x 25 x 1 mm) with red silicone sheets (Diversified Silicone Products, #5038GP-032, 50 Duro, 1/32 in thick; McMaster-Carr #1460N21) in between the glass slides. The flow chamber contains five 8 mm diameter circles spaced 10 mm apart that are connected by a 5mm wide channel (Fig. 1). The silicon sheet is bonded to the bottom glass microscope slide using plasma corona treatment. To visualize and quantify the growth of bacterial colonies in response to interstitial advective fluxes, we created hydrogel disks in the shape of a disk with a 3 mm diameter and 1 mm height and a single 400 µm channel through the particles where we control the advective flux through the particle by changing the angle of the channel relative to the fluid flow. We prepared the hydrogel disks using low-melting agarose as previously described^13^. In brief, we dissolved 15 mg of low melting agarose (Agarose low gelling temperature, Sigma-Aldrich, CAS# 39346-81-1) in 1 mL of LB medium in a block heater at 70 °C. We then cooled the agarose directly in the block heater to 40 °C. Once cooled, we then 10 µl of a bacterial suspension containing 10^7^ cells mL^−1^ (final bacterial concentration in the agar of 10^5^ cells mL^−1^). We then create a 1 mm thick sheet of agarose using a 3 mL syringe (BD, 3 mL Luer lock) and a blunt needle (Industrial Dispensing Supplies, 22G) by dispensing the molten agarose in between two microscope slides that are separated by 1 mm using a separate microscope slide. Once solidified (15 minutes at room temperature, ∼20 °C), we punched the agarose discs from the agarose slab using a 3 mm diameter biopsy punch (Integra Miltex, Disposable Biopsy Punch, 3 mm) and bisected the particles into equal halves using an X-ACTO knife. We then transfer the two halves into the flow chamber and manually positioned the two halves at approximately 400 µm apart with the required angle relative to the fluid flow. Finally, we placed the microscope cover glass containing the inlet and outlet ports on top of the assembly and sealed the whole experimental system by applying pressure to the silicone layer, and reinforced the seal with labeling tape.

### Millifluidic growth conditions and image acquisition

We controlled the flow within the millifluidic device using a syringe pump (Harvard Apparatus, PHD ULTRA). We placed five particles in each millifluidic device with the intra hydrogel disks channel positioned between 0° (inline) and 90° (perpendicular) to the flow to control the relative contribution of advective and diffusive fluxes to the overall mass transport without changing the geometry of the microenvironment. We then imposed a settling velocity of 10 m d^−1^ around the particles (LB media) for 24 hours. After this time, we visualize the bacterial colonies within the particles using a Nikon Ti2-E inverted microscope equipped with an Andor Zyla 4.2 sCMOS camera. The microscope was controlled using Nikon Elements (v.4) and all images were captured in 16-bit with a 10x objective (Plan Fluor DLL). Each hydrogel disk was visualized at five depths 50 µm apart and the center was located approximately at the middle of the channel thickness (i.e. ∼500 µm from the bottom glass slide which we located using the perfect focus system of the Nikon TI2 microscope). Each depth layer was captured as a 4 x 4 tile scan with a 20% overlap that was automatically stitched by the Nikon Elements software. Individual images of the tile scans were captured at an Illuminator Iris intensity of 30 and shutter speed of 50 ms for all particles to ensure equal illumination of each particle and thus reliably quantify the colonized population. These settings resulted in an optimal dynamic range within the images for particles and thus required the least amount of post-processing.

### Digital image analysis and quantification of bacterial colony size

We analyzed the images using MATLAB (R2020) with the image analysis toolbox. Initially, we classify particles based on the presence or absence of a channel using a custom algorithm. This algorithm employed a shape index filter to classify particles based on the measured circularity and elongation shape factor. The latter corresponds to the square root of the ratio of the second moments of an object around its principal axes^42^. Particles that deviated from the expected shape of intact particles were then either classified as half particles or discarded if did not meet half particles characteristics. For the particles matching the geometric characteristics of particles cut in half, they were further processed to extract relevant features of the inner channel (i.e., channel angle and width). We transformed the particles into circles with no channel and then superimposed a second image with the original channel to extract the channel itself following a black hat transform (differences between the closed image and the image itself) following a white hat transform (difference between the input image and its opening). We subsequently utilized this channel to measure the exact angle orientation, width, and length of the channel. Finally, we segmented the colonies and their spatial positions relative to the channel and particle edges were obtained using methods described elsewhere^7,13^.

### Digital twin simulations using the COMSOL simulation software

We used COMSOL Multiphysics v5.6 to create digital twin simulations of the experimental system to observe the spatiotemporal evolution of oxygen and associated bacterial patterns. The simulated geometry and equations used for describing bacterial growth are outlined below.

*Simulation Geometry*.

We created a 2D time-dependent model that contains both the particle and surrounding flow channel (Fig. 1). We used a digital twin approach (i.e., 3 mm diameter particle inside 8 mm diameter chambers that is connected via 5 mm wide channels) and included no-slip boundaries for the flow chamber and particle surface. We include channels ranging from 200 µm to 600 µm inside the particles that are rotated from 0° to 90° in 15° increments with respect to the flow. Due to our inability to precisely control the width and rotation in the experiments, we explore the influence of these two parameters beyond the experimental geometry using the model. We simulate flow fluid flow inside the flow chamber by imposing a mean flow velocity across the whole flow channel (11.5 µm s^−1^, 115 µm s^−1^, and 1150 µm s^-^) which generates flow inside the channel due to the geometry and resulting pressure distribution. In addition to flow through the channels, hydrogels permit flow through the hydrogel matrix due to their nanoporous structure which we can calculate for solid particles as a function of their size and permeability^41^. However, due to the low permeability of hydrogels (e.g., 450 nm^2^ for 3% agarose hydrogel^43^), the fluid collection efficiency is merely 10^−7^% and negligible when compared to the fluid collection efficiency driven by flow through the channels (>10^−2^% for the majority of conditions, Fig. 4b). In addition, since the majority of this flow through the hydrogel is confined to a thin boundary layer^41^ (in our case ∼50 nm assuming a linear relationship between agarose concentration and permeability), elevated nutrient supply due to flow through the hydrogel matrix only affects regions where diffusive fluxes dominate (characteristic time of diffusion for oxygen across 50 nm is on the order of 1 µs) and thus do not significantly alter nutrient fluxes. For this reason, we simulated the particles as impermeable solids in COMSOL where nutrient fluxes inside the particles are limited to diffusion. Finally, we used the COMSOL standard diffusion coefficient for oxygen in the water and agarose of 1.0*10^−9^ m^2^ s^−1^ and assigned a cell density of 10^5^ cells mL^−1^ inside the particle with negligible diffusion of the cells to simulate immobilized cells. We set the oxygen concentration at the inflow and boundary of the flow chamber (silicone) as fully saturated (258 µmol L^−1^).

*Bacterial growth and oxygen consumption*.

Bacterial growth was simulated using a population-based approach with a Monod-type oxygen limitation term to connect the underlying oxygen distributions to the emerging bacterial growth patterns. We additionally limited bacterial growth using a carrying capacity term to avoid physically impossible bacterial densities (exceeding space-filling conditions). We simulated the bacterial cells as a concentration of cells and represented the oxygen consumption and bacterial growth as reaction terms as shown in Equations 2 and 3.

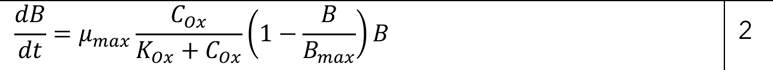

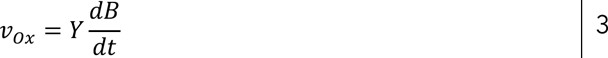

Here, B is the concentration of cells [cells m^−3^], µ_max_ the maximum growth rate [s^−1^], C_ox_ the oxygen concentration [mM], K_ox_ is the oxygen half-saturation coefficient [mM], B_max_ is the carrying capacity that translates to the whole space filled by 1 µm^3^ bacterial cells [10^18^ cells m^−3^], *v_ox_* is the oxygen consumption rate [mol cell^−1^ s^−1^] and Y the oxygen demand [mol cell^−1^]. Values for all parameters are shown in Table 1. In addition, we ensured non-negative oxygen and cell densities for solver stability.

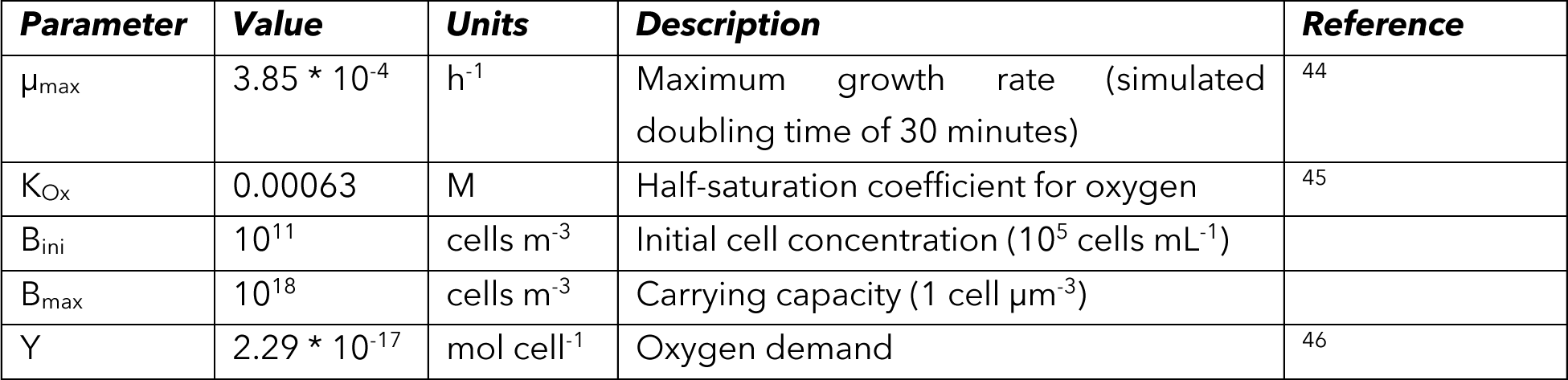

## Author contributions

BB and ARB developed the research question and designed the experiments. RS and BB performed the experiments and COMSOL simulations. DC analyzed the microscopy images. BB analyzed the model and experimental data and performed statistical tests. All authors contributed to writing the manuscript and approved the final version of the manuscript.

## Acknowledgments

This study was funded by the National Science Foundation OCE-2142998 and the Simons Foundation grant number #622065 awarded to ARB. RS was funded partially through the MIT UROP Office and the MIT NEET Program provided access to COMSOL Multiphysics for much of the computational modeling work. BB was supported by a fellowship of the Swiss National Science Foundation grant number P500PN_202842. The funders played no role in the study design, data collection, analysis, and interpretation of data, or the writing of this manuscript.

## Data and code availability

All extracted data and the underlying code for this study are available in Zenodo and can be accessed via this DOI: 10.5281/zenodo.8338298.

