## Supplementary Information for "Physical structure and interstitial flows govern microbial life in microenvironments"

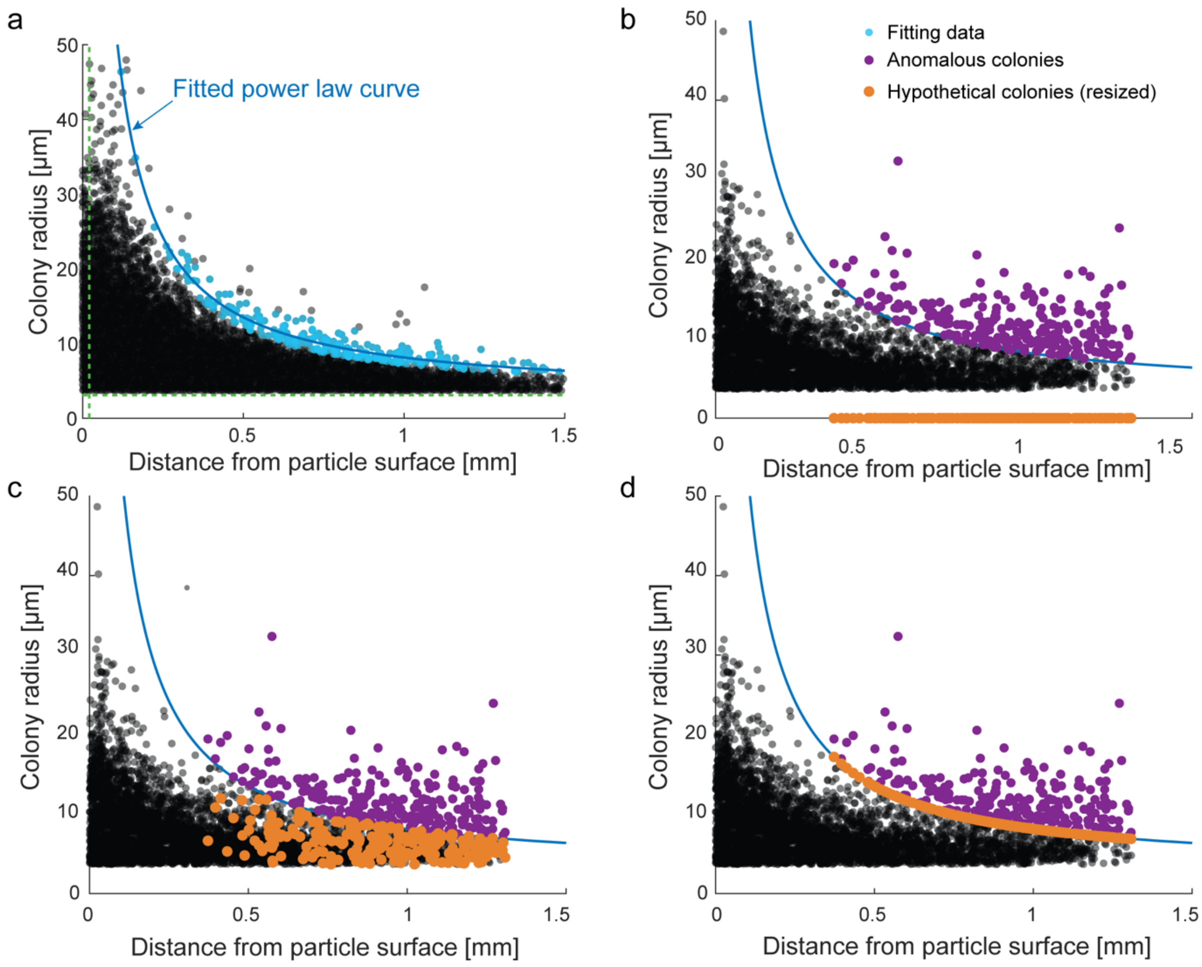


**Supplementary Figure 1: Visualization of the approach used to determine the relative biomass increase. a)** We fitted a modified power law equation (see methods section) to the envelope of the colony size versus distance from the particle periphery diagram (fitting data shown in blue). We obtained the fitting data by using the same modified power law equation with initial estimates based on physical properties from the experimental system (particle boundary thickness of 50 µm for the x-axis shift and minimum colony size for the y-axis shift, both denoted by dotted green lines) and selected colonies that deviate less than 2 µm from this hypothetical line. We then refitted the modified power law to the fitting data to obtain the final curve which we used to determine anomalous colonies in particles with a channel. We used three approaches to determine the relative biomass increase due to the presence of advective fluxes by calculating the ratio of the total observed biomass (including the anomalous colonies) and hypothetical biomass by omitting the anomalous colonies (**b**), assigning a colony radius between the minimum radius and fitted curve (**c**) or assigning a radius equal to the fitted curve (**d**).

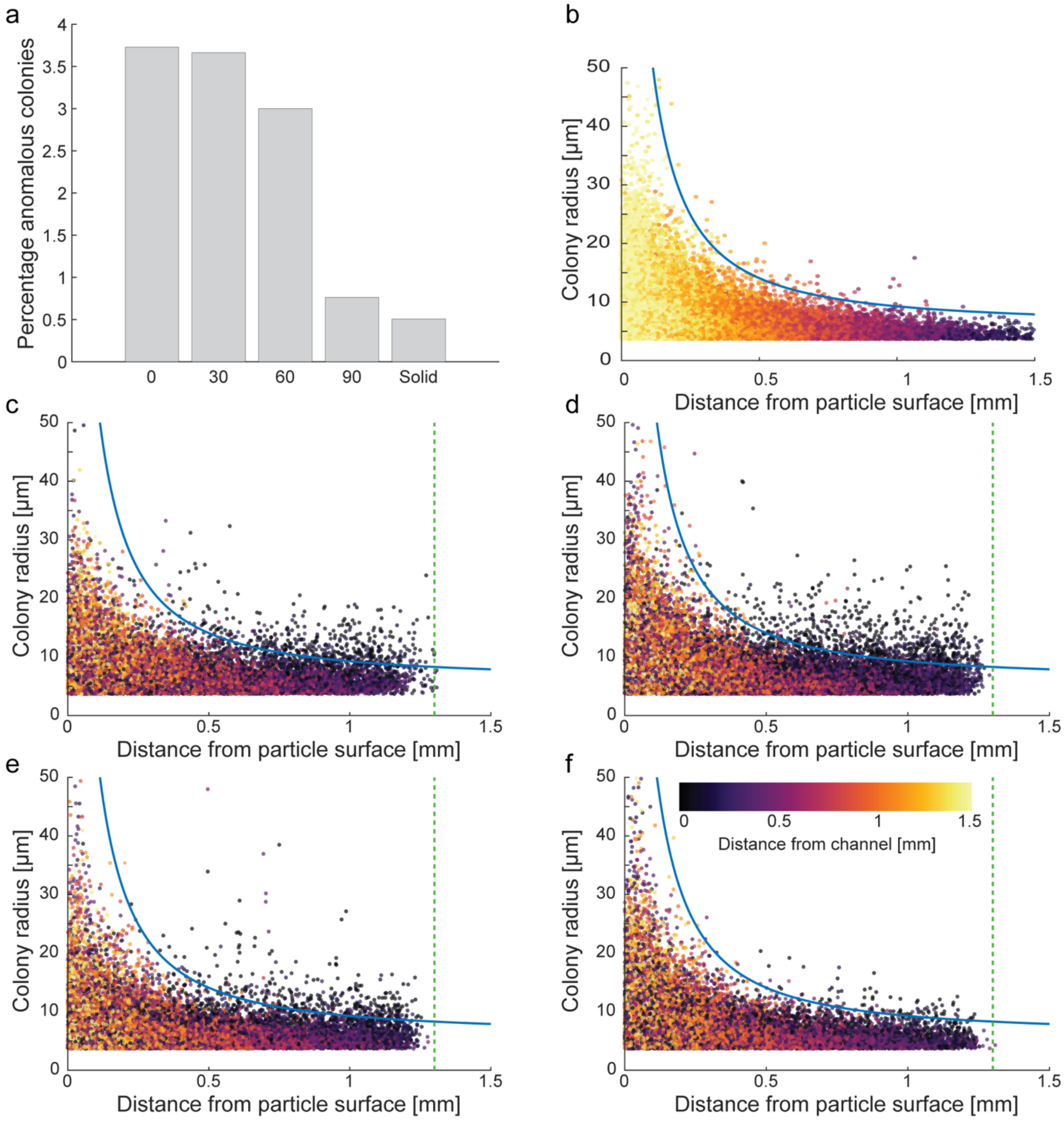


**Supplementary Figure 2: Quantification and visualization of anomalous colonies in particles with a channel. a)** Mean percentage of anomalous colonies identified in particles with different channel angles. **b)** Colony radius versus distance from the particle surface for the solid particles. Since these particles do not have a channel, the color indicates the distance from the particle center. Colony radius versus distance from the particle surface for particles with a channel angle of 0° (**c**), channel angle of 30° (**d**), channel angle of 60° (**e**), and channel angle of 90° (**f**). Typically, anomalous colonies (colonies above the blue threshold) are far from the particle surface but close to the channel as indicated by the color scheme.

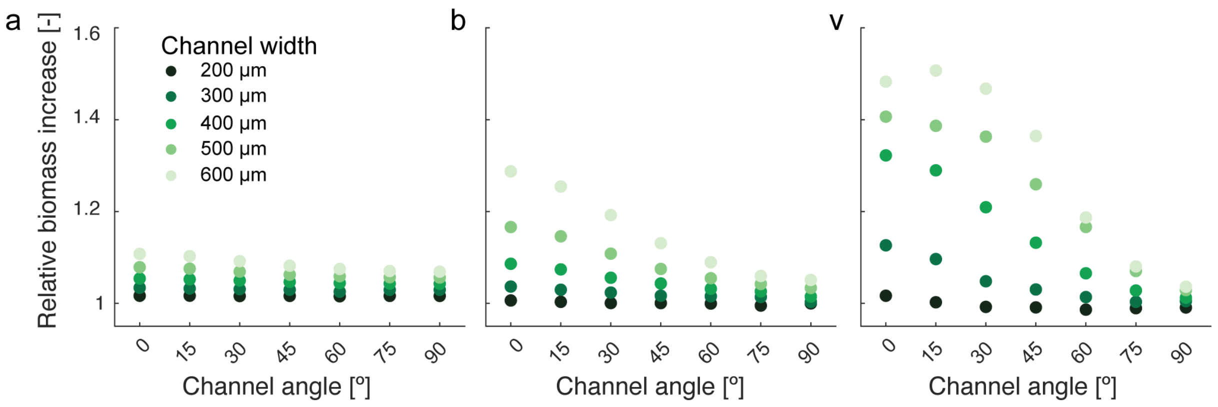


**Supplementary Figure 3: Relative biomass increase as a function of flow velocity.** We use COMSOL to characterize the influence of the flow velocity on the relative biomass increase by simulating a flow velocity of 11.5 µm $s^{-1}$ (**a**), congruent flow velocity of 115 µm $s^{-1}$ (**b**) or increasing to 1150 µm $s^{-1}$ (**c**). The data underlying this figure is provided in Supplementary Table 2.

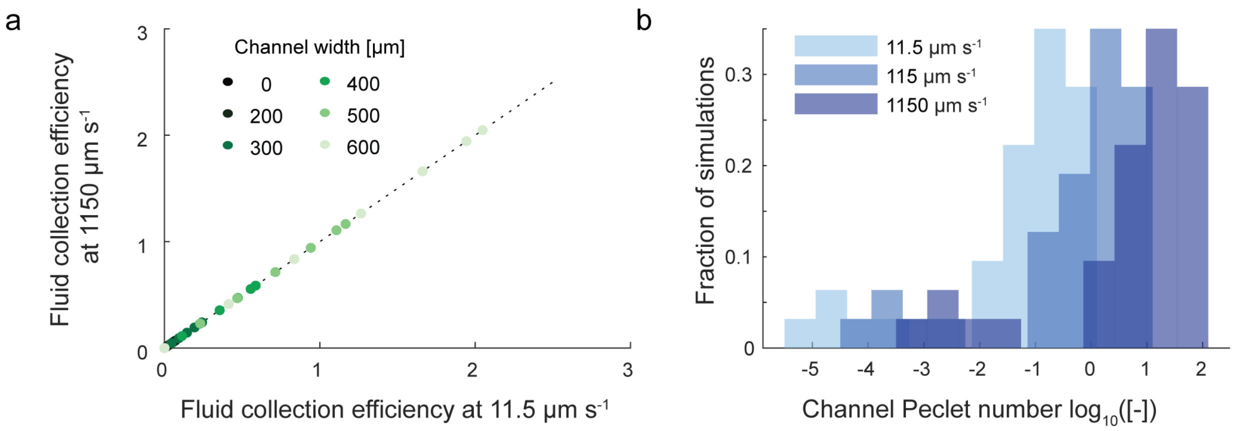


**Supplementary Figure 4: Influence of flow velocity on the fluid collection efficiency and channel Pe. a**) The fluid collection efficiency is not affected by the flow velocity as shown between the comparison at 11.5 µm $s^{-1}$ flow velocity and 1150 µm $s^{-1}$ flow velocity. **b**) The flow velocity shifts the magnitude of Pe associated with the flow and geometry of the channel, but not the pattern associated with different geometries.

**Supplementary Table 1: Relative biomass increase for all channel angles depending on the analysis algorithm.** Relative biomass increase is calculated by omitting large colonies, stochastically reducing large colonies to a size between 0 and the upper limit of the fitted power law and reducing to the characteristic size determined by the fitted power law curve. Differences in the analysis procedure are highlighted in Supplementary Fig. 2. Values are given as mean ± *SD*.

| | **0°** | **30°** | **60°** | **90°** | **Solid** |
|---|---|---|---|---|---|
| **Omitted** | *7.52% ± 4.08%* | *7.39% ± 4.55%* | *5.82% ± 2.96%* | *1.18% ± 1.33%* | *1.38% ± 1.45%* |
| **Stochastic** | *5.16% ± 2.90%* | *5.25% ± 3.32%* | *4.09% ± 2.24%* | *0.73% ± 0.85%* | *0.80% ± 0.85%* |
| **Reduced** | *2.93% ± 1.85%* | *3.07% ± 2.10%* | *2.33% ± 1.39%* | *0.33% ± 0.42%* | *0.34% ± 0.38%* |

**Supplementary Table 2: Percentage biomass increase for all channel angles and widths depending on the bulk flow velocity.** We attribute negative values of relative biomass increase at medium and high flow velocities to small deviations in the flow field and numerical error.

| | | ***Channel width*** | | | | | | | | | | | | | | |
|---|---|---|---|---|---|---|---|---|---|---|---|---|---|---|---|---|
| | | *200 µm* | | | *300 µm* | | | *400 µm* | | | *500 µm* | | | *600 µm* | | |
| *Bulk velocity (µm $s^{-1}$)* | | *11.5* | *115* | *1150* | *11.5* | *115* | *1150* | 11.5 | 115 | 1150 | 11.5 | 115 | 1150 | 11.5 | 115 | 1150 |
| ***Channel angle*** | *0°* | 1.6 | 0.6 | 1.7 | 3.4 | 3.7 | 12.6 | 5.4 | 8.6 | 32.2 | 7.9 | 16.6 | 40.7 | 10.8 | 28.8 | 48.3 |
| | *15°* | 1.7 | 0.3 | 0.2 | 3.2 | 3.0 | 9.6 | 5.2 | 7.4 | 29.0 | 7.6 | 14.6 | 38.7 | 10.3 | 25.4 | 50.7 |
| | *30°* | 1.6 | 0.1 | -0.8 | 3.1 | 2.3 | 4.8 | 5.0 | 5.6 | 21.0 | 6.9 | 10.8 | 36.3 | 9.2 | 19.2 | 46.8 |
| | *45°* | 1.6 | 0.1 | -0.9 | 3.0 | 1.7 | 3.0 | 4.7 | 4.3 | 13.2 | 6.2 | 7.5 | 25.9 | 8.2 | 13.1 | 36.5 |
| | *60°* | 1.5 | 0.0 | -1.4 | 2.5 | 1.5 | 1.4 | 4.4 | 3.2 | 6.6 | 5.9 | 5.5 | 16.6 | 7.5 | 8.9 | 18.7 |
| | *75°* | 1.6 | -0.5 | -1.0 | 2.9 | 1.4 | 0.3 | 4.3 | 2.6 | 2.8 | 5.7 | 4.2 | 7.0 | 7.0 | 6.0 | 8.0 |
| | *90°* | 1.6 | 0.0 | -1.0 | 3.0 | 0.3 | 0.5 | 4.3 | 1.5 | 1.2 | 5.6 | 3.3 | 2.9 | 6.9 | 5.1 | 3.6 |

**Supplementary Table 3: Maximum oxygen penetration for all channel angles and widths depending on the bulk flow velocity.** Oxygen penetration is given in percentage of the channel length (3 mm).

| | | ***Channel width*** | | | | | | | | | | | | | | |
|---|---|---|---|---|---|---|---|---|---|---|---|---|---|---|---|---|
| | | *200 µm* | | | *300 µm* | | | *400 µm* | | | *500 µm* | | | *600 µm* | | |
| *Bulk velocity (µm $s^{-1}$)* | | *11.5* | *115* | *1150* | *11.5* | *115* | *1150* | 11.5 | 115 | 1150 | 11.5 | 115 | 1150 | 11.5 | 115 | 1150 |
| ***Channel angle*** | *0°* | 6 | 8 | 18 | 11 | 17 | 83 | 17 | 36 | 100 | 25 | 72 | 100 | 34 | 100 | 100 |
| | *15°* | 6 | 8 | 18 | 11 | 17 | 76 | 17 | 33 | 100 | 25 | 69 | 100 | 34 | 100 | 100 |
| | *30°* | 6 | 8 | 16 | 11 | 15 | 64 | 16 | 29 | 100 | 23 | 59 | 100 | 31 | 100 | 100 |
| | *45°* | 6 | 8 | 12 | 11 | 14 | 47 | 15 | 25 | 100 | 23 | 48 | 100 | 28 | 86 | 100 |
| | *60°* | 6 | 7 | 11 | 10 | 13 | 33 | 15 | 22 | 90 | 20 | 38 | 100 | 28 | 62 | 100 |
| | *75°* | 6 | 6 | 8 | 10 | 11 | 19 | 15 | 17 | 46 | 20 | 27 | 100 | 25 | 41 | 100 |
| | *90°* | 6 | 6 | 6 | 10 | 11 | 11 | 14 | 15 | 15 | 18 | 20 | 20 | 23 | 26 | 26 |

**Supplementary Table 4: Maximum channel velocity for all channel angles and widths depending on the bulk flow velocity.** Pore velocities are extracted from the simulations at the center of the pore (i.e., equidistant from both the channel walls and entrances).

| | | ***Channel width*** | | | | | | | | | | | | | | |
|---|---|---|---|---|---|---|---|---|---|---|---|---|---|---|---|---|
| | | *200 μm* | | | *300 μm* | | | *400 μm* | | | *500 μm* | | | *600 μm* | | |
| *Bulk velocity ($\mu m\ s^{-1}$)* | | *11.5* | *115* | *1150* | *11.5* | *115* | *1150* | 11.5 | 115 | 1150 | 11.5 | 115 | 1150 | 11.5 | 115 | 1150 |
| ***Channel angle*** | *0°* | 0.2 | 2.1 | 21.1 | 0.5 | 4.9 | 48.7 | 0.7 | 8.6 | 86.4 | 1.4 | 13.9 | 138.1 | 2.0 | 20.0 | 199.7 |
| | *15°* | 0.2 | 2.0 | 20.0 | 0.5 | 4.6 | 46.1 | 0.8 | 8.2 | 81.7 | 1.3 | 13.1 | 131.0 | 1.9 | 19.0 | 190.1 |
| | *30°* | 0.2 | 1.3 | 16.9 | 0.4 | 3.9 | 38.9 | 0.7 | 7.0 | 69.7 | 1.1 | 11.1 | 111.4 | 1.6 | 16.2 | 161.9 |
| | *45°* | 0.1 | 1.2 | 12.6 | 0.3 | 2.9 | 29.1 | 0.5 | 5.2 | 52.4 | 0.8 | 8.4 | 84.3 | 1.2 | 12.3 | 122.8 |
| | *60°* | 0.1 | 0.8 | 8.1 | 0.2 | 1.9 | 18.9 | 0.3 | 3.4 | 33.9 | 0.6 | 5.5 | 55.4 | 0.8 | 8.2 | 81.7 |
| | *75°* | 0.0 | 0.4 | 4.0 | 0.1 | 0.9 | 9.3 | 0.2 | 1.7 | 16.8 | 0.3 | 2.7 | 27.3 | 0.4 | 4.0 | 40.2 |
| | *90°* | 0.0 | 0.0 | 0.0 | 0.0 | 0.0 | 0.0 | 0.0 | 0.0 | 0.0 | 0.0 | 0.0 | 0.0 | 0.0 | 0.0 | 0.0 |
